# O-linked mucin-type glycosylation regulates the transcriptional programme downstream of EGFR in breast cancer

**DOI:** 10.1101/714675

**Authors:** Virginia Tajadura-Ortega, Gennaro Gambardella, Alexandra Skinner, Katrine Ter-Borch Gram Schjoldager, Richard Beatson, Rosalind Graham, Daniela Achkova, Joyce Taylor-Papadimitriou, Francesca D. Ciccarelli, Joy M. Burchell

## Abstract

Aberrant mucin type O-linked glycosylation is a common occurrence in cancer. This type of O-linked glycosylation is not limited to mucins but can occur on many cell surface glycoproteins where only a small number of sites may be present. Upon EGF ligation, EGFR induces a signaling cascade but can also translocate to the nucleus where it can directly regulate gene transcription. Here we show that upon EGF binding, human breast cancer cells carrying different O-linked glycans respond by transcribing different gene expression signatures. This is not a result of changes in signal transduction but due to the differential nuclear translocation of EGFR in the two glyco-phenotypes. This is regulated by the formation of an EGFR/galectin-3/MUC1/β-catenin complex at the cell surface that is present in cells carrying short core-1-based O-glycans characteristic of tumour cells but absent in core-2-carrying cells.

## INTRODUCTION

Glycosylation of proteins is the most abundant post-translational modification and greatly increases the size, diversity and function of the proteome. Nearly all proteins that are expressed on the cell membrane or are secreted carry glycans that are involved in cell adhesion, recognition, molecular trafficking, clearance and signaling (1). Moreover, aberrant glycosylation occurs in essentially all types of human cancer making this one of the hallmarks of malignancy. Indeed, changes in glycosylation appear to be early events, as well as playing key roles in the induction of invasion and metastases (2, 3).

O-linked mucin-type glycosylation (here referred to as O-linked glycosylation) is characterized by the addition of GalNAc to serine or threonine residues of proteins. Although found abundantly on mucins, which are heavily O-linked glycosylated, this type of glycosylation is also found on many other types of glycoproteins including those involved in signal transduction (4, 5). Changes in O-linked glycosylation is one of the most prevalent glyco-phenotypes observed in solid tumors (6) and often results in O-linked glycoproteins carrying short, linear sialylated glycans rather than the longer branched glycans seen in normal epithelial cells (7). However, the presence of short linear or branched chains is not mutually exclusive and particularly in estrogen receptor negative breast cancers the O-linked glycans can contain branched glycans (8) (figure S1a). Aberrant O-glycosylation promotes tumour growth (9, 10), leads to remodeling of the microenvironment (11) and increases metastatic potential (8, 12).

Members of the epidermal growth factor receptor family (ErbBs) play a major role in cancer. ErbB2 (HER2) is involved in driving tumorigenesis in breast cancer and ErbB1 (EFGR) signaling is frequently dysregulated in cancer. Overexpression and mutation of EGFR is seen in many solid tumours. In addition to HER2, EGFR is an important therapeutic target with a number of small molecular inhibitors and anti-EGFR antibodies being in clinical use (13). EGFR is over-expressed in triple negative breast cancers but activation of its signaling pathway is important in other sub-types of breast cancer (14).

A number of proteins have been shown to interact with EGFR including the highly O-glycosylated mucin, MUC1, which is upregulated and aberrantly glycosylated in breast and other carcinomas. This interaction is reported to modulate the stability of EGFR by preventing its ubiquitination. This results in the recycling of EGFR rather than its degradation (15). In turn, EGFR can phosphorylate the cytoplasmic tail of MUC1 (CT-MUC1) (16). Moreover, CT-MUC1 interacts with EGFR in the cell nucleus and promotes the binding of EGFR to chromatin and the *CCND1* (cyclin D1) promoter (17).

Given the interaction of EGFR with MUC1, we investigated the effects of different O-linked glycosylation patterns on the transcriptional program promoted by EGF in breast carcinoma cells. We show that different glycosylation patterns of the cell lead to the differential expression of proteins induced after EGF binding, showing for the first time that the O-linked glycosylation pattern of the cell plays a role in controlling EGF induced gene expression. These results give an additional insight into the role of aberrant O-linked glycosylation in tumorigenesis.

## RESULTS

### The gene expression profile of cells treated with EGF changes with alterations in cellular O-glycosylation

To determine if O-linked glycosylation influences the transcription programme induced upon stimulation with EGF, we used the breast carcinoma cell line T47D that expresses short, linear O-glycans and isogenic cell lines engineered to express branched core 2 O-glycans through the transfection of *GCNT1* encoding the C2GnT1 glycosyltransferase (fig S1a), and which we refer to as T47D-C2GnT1 (18). We isolated two independent clones T47D-C2GnT1-J and T47D-C2GnT1-B which overexpressed *GCNT1* by at least 30 fold (fig S1b). All the lines expressed MUC1 as shown by staining with the HMFG1 antibody (fig S1c). However, both T47D-C2GnT1-J and T47D-C2GnT1-B cells had reduced staining with the anti-MUC1 monoclonal antibody SM3 (fig S1c). As the binding of SM3 to its epitope within MUC1 is inhibited by core 2 glycans (9, 18) this confirms that glycoproteins expressed by T47D-C2GnT1-J and -B carry branched core 2 glycans. We also examined the surface expression of EGFR by flow cytometry in T47D cells and the two different T47D-C2GnT1 clones and observed that all three lines had similar percentage of cells expressing EGFR (fig S1c).

We investigated the expression of Cyclin D1, one of the known transcriptional targets of EGFR (19) in T47D and T47D-C2GnT1-J cells stimulated with EGF for several time intervals. We observed that the expression of the Cyclin D1 mRNA (fig 1a) and protein (fig 1b) peaked after 6h of treatment with EGF in T47D carrying either short, sialylated core 1-based O-glycans (T47D) or branched core 2-based O-glycans (T47D-C2GnT1-J). However, a significantly higher level of expression was observed in T47D-C2GnT1-J cells after 6h of treatment with EGF (fig 1a-b).

**Figure 1.**
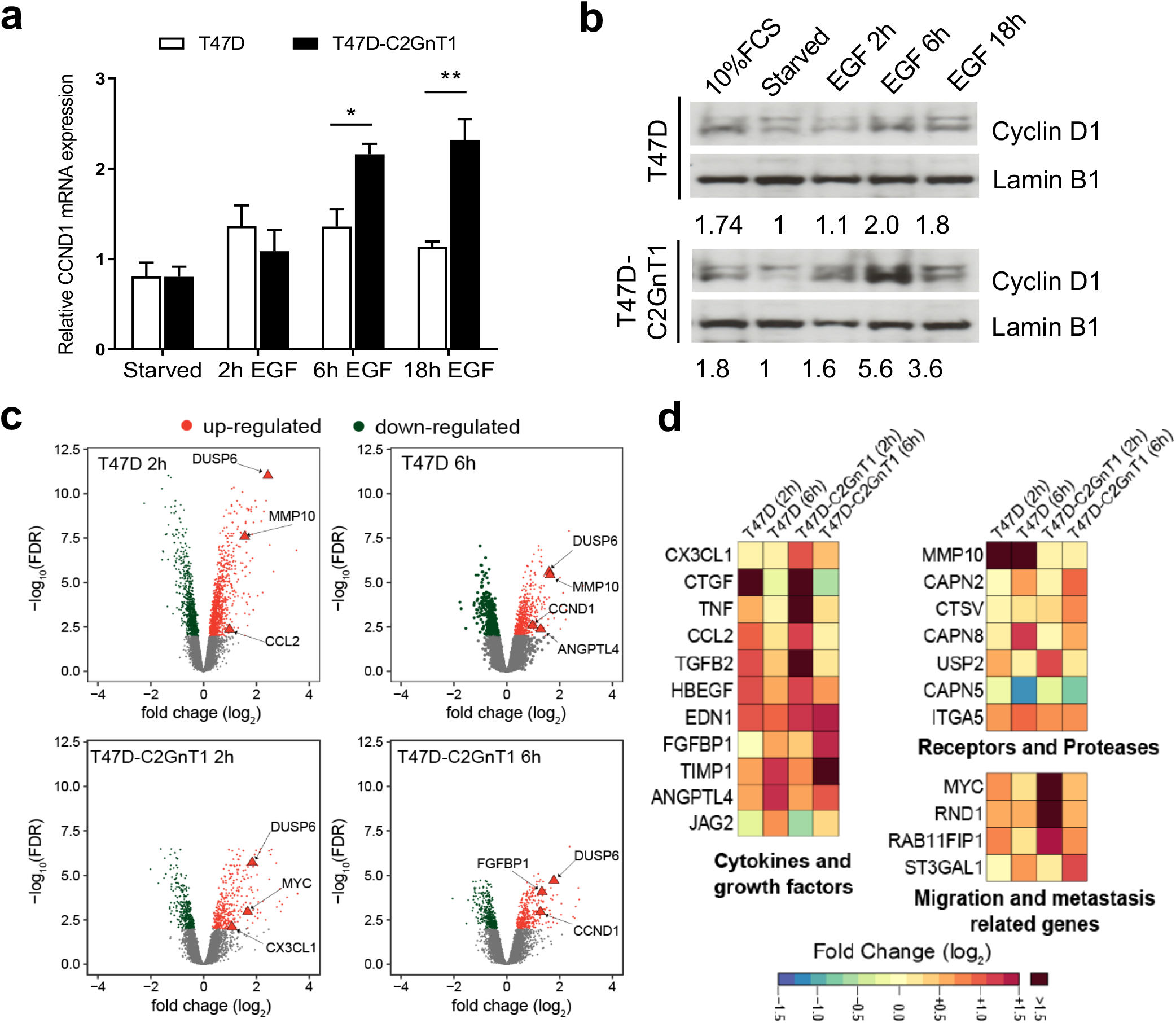
EGF induced the differential expression of microenvironment modulating factors in cells with different O-glycosylation. T47D and T47D-C2GnT1-J cells were starved for 24 hours and treated with EGF (100ng/ml) for the indicated times or 10% FCS for 2 h. (a) Transcript levels of Cyclin D1 were assessed by qPCR using PUM1 as a housekeeping gene, quantified using the DDCt method (**P<0.01, ***P<0.001; analysis by t-student test n=4). (b) Cell extracts were subjected to western blotting using the indicated antibodies. Representative western blots are shown. Figures report the densitometric ratio of Cyclin D1 to Lamin B1. (c) Analysis of RNA expression using an Illumina Human HT12 v4 beadchip. Volcano plots show mean fold change (log2) in gene expression on x axis and −log10(FDR) on y axis. Significantly differentially expressed genes (FDR < 0.01) after 2h or 6h of treatment with EGF are highlighted in red (up-regulated) or green (down-regulated). Relevant transcriptional targets of EGF signaling are highlighted as triangles. (d) Expression matrix of genes belonging to the cytokines and growth factors, receptors and proteases and migration and metastasis related genes categories. Selected genes whose expression was induced to at least twice the baseline (log2 FC>1) in at least one condition are shown. Genes with a log2 FC>1.5 are represented with the same colour.

To study the transcriptional program induced upon stimulation with EGF in cells with different O-glycosylation patterns, genome wide expression microarrays were performed with T47D and T47D-C2GnT1-J cells stimulated with EGF for 2h and 6h. Upon EGF stimulation, approximately 8% of the transcripts where significantly modulated (FDR < 0.01) after 2h or 6h of treatment in either T47D or T47D-C2GnT1-J cells (fig 1c and table S1). As expected Cyclin D1 (*CCND1*) was confirmed to be up-regulated, with twice the expression of the baseline (log_2_ fold change >= 1, fig 1c) after 6h of treatment. Some transcriptional targets of EGFR like the DUSP6 phosphatase that is transcriptionally induced by EGF signaling (20, 21) was found to be upregulated in both cell lines at both time points (fig 1c). However, we found that some genes (such as MMP10 or ANGPTL4) were overexpressed in T47D but not in T47D-C2GnT1-J, while others like FGFBP1, CX3CL1 and MYC were more highly expressed in T47D-C2GnT1-J (fig 1c).

Gene ontology enrichment analysis (GOEA) (table S2) of overexpressed genes with at least two fold change (log_2_ fold change >= 1) as compared to non-stimulated conditions confirmed a strong enrichment for transcription factor activity and positive regulation of protein phosphorylation as was described before for EGF stimulation of breast primary and cancer cells in (21). We selected a subset of 20 genes that showed differential expression between the two glyco-phenotypes for further validation. These were classified into 4 categories: (i) membrane receptors and proteases (ii) cytokines and growth factors (iii) migration and metastasis related genes and (iv) transcription factors (fig 1d). We confirmed the differential expression between T47D and T47D-C2GnT1-J of 16 out of the 20 by quantitative real-time PCR (table S3). We also performed a global comparison of the difference in gene expression between the paired cell lines T47D and T47D-C2GnT1-J at 0, 2 and 6 hours after EGF stimulation (fig S2a and table S1). Among the genes differentally expressed between the two cell lines at baseline, 2 or 6 hours of treatment with EGF, we identified most of the genes validated previously (fig 1c-d and table S3). These data show that the O-linked glycosylation status of the cell had a significant effect on the EGF induced transcriptional profile of the cells, in particular we found that the O-linked glycosylation affected the level of expression induced by EGF stimulation with many genes showing enhanced expression in T47D-C2GnT1 cells compared to T47D cells.

Finally, we confirmed the expression of three differentially expressed genes (*FGF-BP1*, *CX3CL1 and MMP10*) in an independent set of samples (fig 2a) and in a second clone of T47D cells transfected with *GCNT1*, T47D-C2GnT1-B (fig S2b) to confirm that the expression of these different transcriptional programs is due to the different O-glycosylation of the cells and not due to differential expression between the clones.

**Figure 2:**
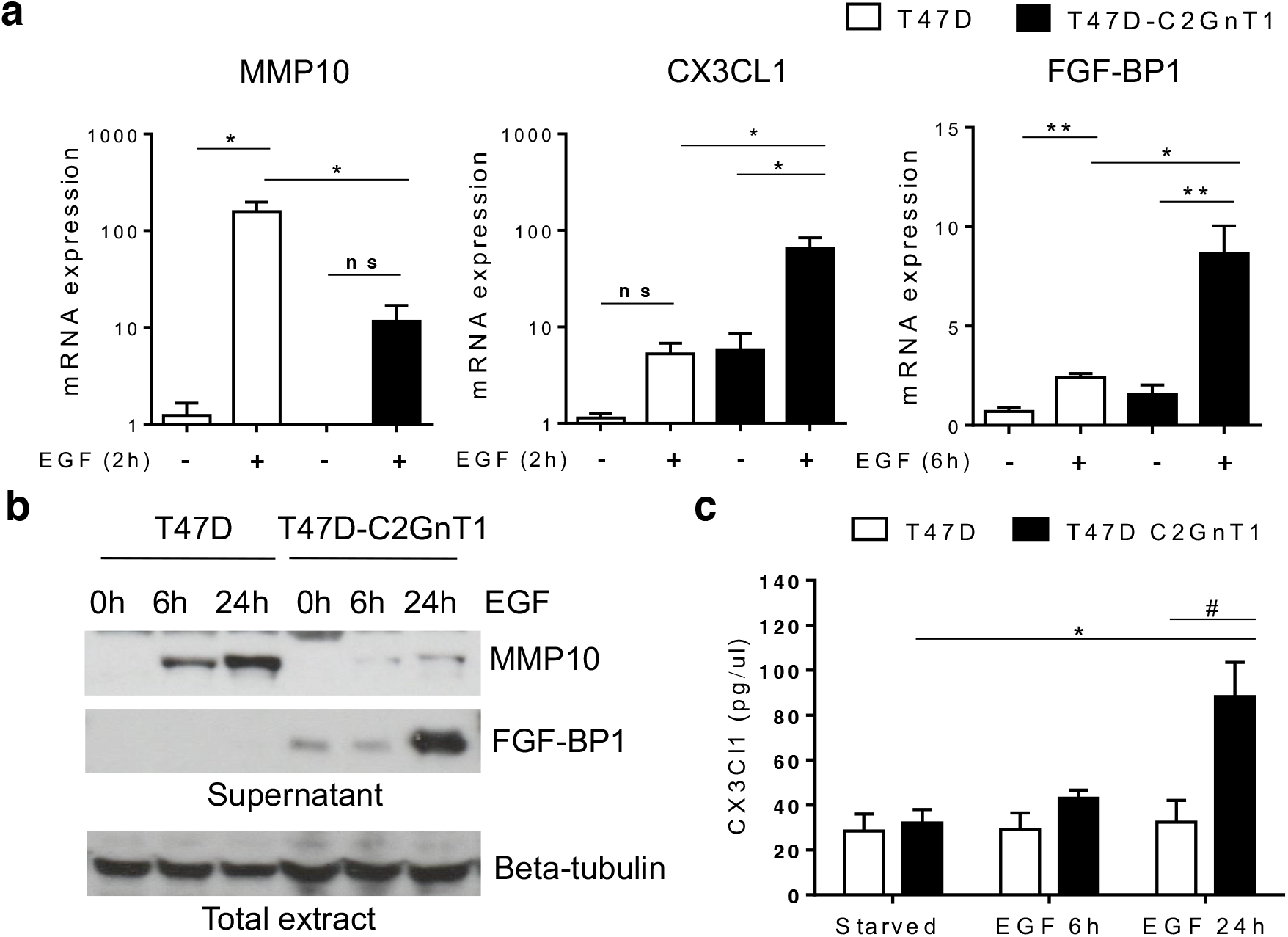
Differential gene expression between the two glycophenotypes results in differential protein expression. T47D and T47D-C2GnT1-J cells were starved for 24 hours and treated with EGF (100ng/ml) for 2h (MMP10 and CX3CL1) or 6h (FGFBP1) (a), or the indicated times (b-c). (a) Transcript levels were assessed by qPCR using PUM1 as a housekeeping gene, quantified using the ΔΔCt method and shown relative to the expression in starved T47D cells (*P<0.05, **P<0.01; analysis by t-student test n=4). (b-c) Total extracts and supernatants were collected and subjected to western blotting with MMP10, FGFBP1 (b) or ELISA to detect CX3CL1 levels (c). (b) Beta-tubulin levels were used as western blot loading control. Representative western blots are shown. (c) CX3CL1 concentration of was measured by ELISA and normalized to the expression of beta-tubulin in total extracts (# P<0.05, T47D vs T47D-C2GnT1-J cells, *P<0.05 vs starved, non-treated conditions; analysis by t-student test n=3).

### EGF induced the differential release of microenvironment modulating factors in cells with different O-glycosylation

As *MMP10*, *CX3CL1* and *FGF-BP1* have all been associated with cancer progression we chose this subset of genes for further study. We observed that the protein expression of the matrix metalloproteinase MMP10 was strongly induced in T47D cells after 6h of EGF stimulation (fig 2b) but was very weakly expressed in T47D-C2GnT1-J cells. In contrast, FGF-BP1 showed no protein induction detectable by immunoblots in the supernatant of T47D cells, but it was significantly induced after EGF stimulation in T47D-C2GnT1-J (fig 2b). Moreover, the chemokine CXC3L1 was increased in the supernatants of EGF stimulated T47D-C2GnT1-J (fig 2c) whereas T47D cells showed no increase in secreted CXC3L1 at 6 hours or 24 hours after EGF stimulation. Together, these data confirm that the differential mRNA expression between T47D and T47D-C2GnT1-J of a subset of genes induced by EGF results in the differential protein expression of these genes by the two glyco-phenotypes.

### CX3CL1, MMP10 and FGF-BP1 expression in breast cancer

In most breast and many other adenocarcinomas the dominant O-linked glycans are linear, short sialylated glycans based on core 1 (fig S1a), whereas in normal breast epithelial cells branched core 2 glycans are exclusively found (7). However, in ER negative (ER-ve) breast cancers core 2 based glycans appear to be the dominant O-linked glycans carried on glycoproteins (8). In a glyco-related gene expression analysis of ER positive (ER+ve) and ER-ve primary breast cancer samples we observed that the ER-ve breast tumours (that carry branched core 2 glycans) tested, showed an increase in CX3CL1 expression (fig 3a) which correlates with our *in vitro* microarray expression data showing increased CX3CL1 expression in the core 2 O-glycan carrying cell line (T47D-C2GnT1) compared with T47D. Analysing the METRABRIC database of 1874 breast cancer samples we found that CX3CL1 and FGF-BP1 were significantly upregulated in ER-ve breast cancers compared to ER+ve (fig 3b. pvalue <1.4×10^−45^ effectively zero)). In contrast, MMP10 was upregulated in ER+ve breast cancer compared to ER-ve (fig 3b). We also observed this correlation with estrogen receptor status in breast cancer cell lines expressing MUC1 and EGFR (data not shown). Our analysis of expression data deposited in Oncomine (from the TCGA breast cancer dataset) shows that MMP10 mRNA is upregulated in breast cancers compared to the normal gland (fig 3c) which exclusively carries branched elongated core 2 glycans (see fig S1a). Since most breast cancers are ER+ve (75%) and those carry predominantly core 1 based glycans, the MMP10 expression in tumours show a correlation with our findings *in vitro*.

**Figure 3:**
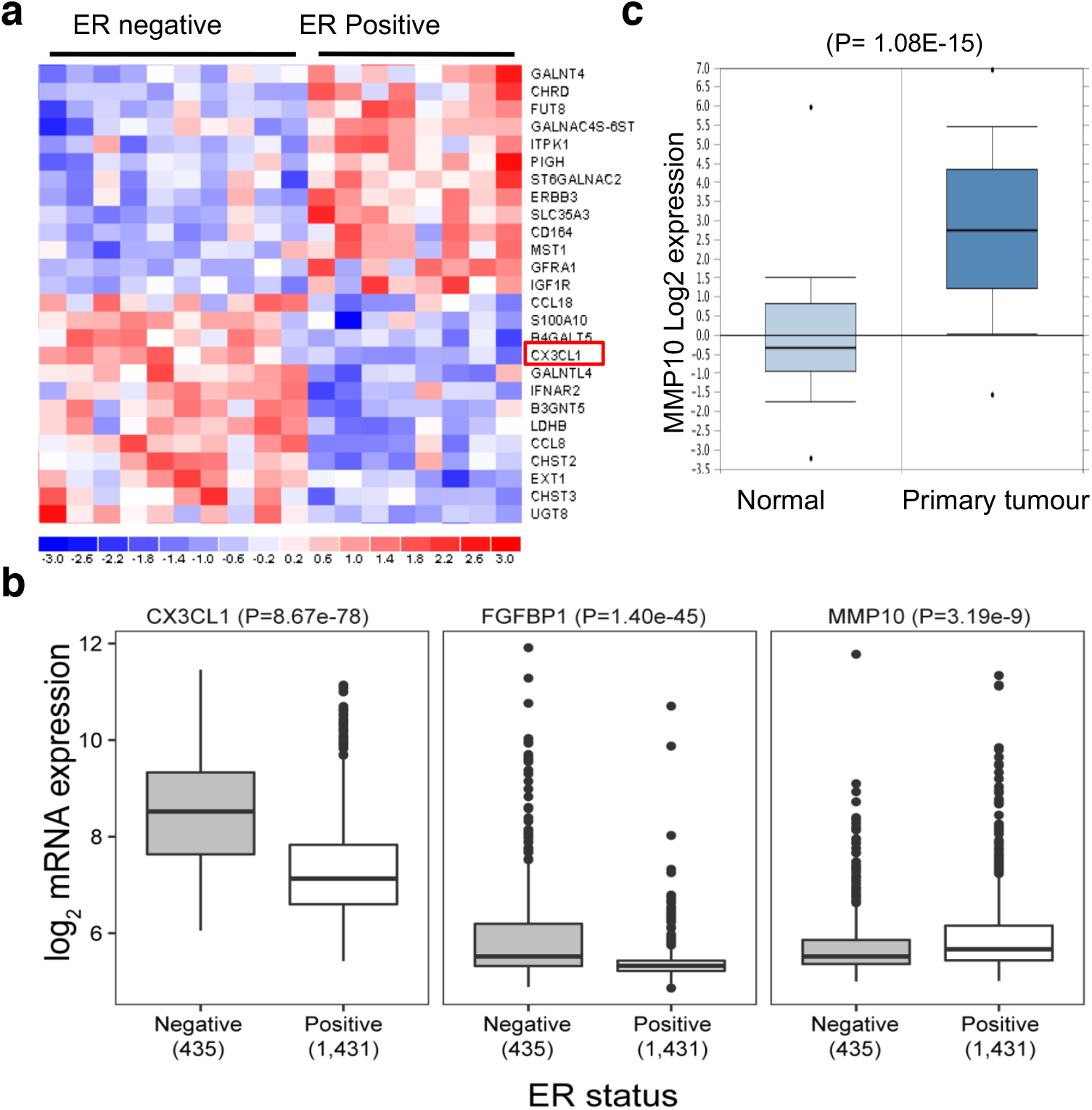
CX3CL1, MMP10 and FGF-BP1 are differentially expressed in ER+ and ER-ve breast cancers. (a) Total RNA was extracted from 10 ER-negative and 8 ER-positive primary breast cancers and submitted to microarray analysis on the GlycoV4 oligonucleotide array shows that CX3CL1 is up-regulated in estrogen receptor negative (ER-ve) primary breast cancer samples. (b) Analysis of 1874 breast cancers in the METABRIC data set for expression of FGF-BP1; CX3CL1 and MMP10, p-values as shown (Mann-Whitney U test). (c) Analysis 76 invasive breast cancer and 61 normal breast samples from The Genome Atlas data set (TCGA) deposited in Oncomine shows upregulation of MMP10 mRNA in breast cancer primary tumours compared to normal breast tissue.

### EGFR accumulation in nuclear endosomes but not activation of EGFR signaling is enhanced in cancer cells carrying core 2 O-glycans

To investigate the mechanisms whereby differences in O-linked glycosylation could influence gene transcription in response to EGF, we looked at EGFR signaling in response to EGF in T47D and T47D-C2GnT-J. The cells were treated with EGF for short time intervals and the activation of EGFR, ERK, AKT and STAT3 downstream signaling pathways examined (fig 4a-b and fig S3a-b). Although total and surface expression of EGFR is similar in both lines in resting conditions (fig S1c and fig 4a-b), we observed a significant difference in the total levels of EGFR between T47D and T47D-C2GnT1-J cells after 10 min of EGF stimulation (fig 4a-b) However, the ratio of phosphorylated EGFR/total EGFR was similar in both cell lines upon EGF stimulation indicating that O-glycosylation does not affect the binding of EGF to its receptor and/or the subsequent autophosphorylation of EGFR (fig 4b). Despite higher levels of EGFR in T47D-C2GnT1-J cells, the activation of downstream ERK and AKT pathways was similar in both cell lines (fig S3a). Moreover, both cell lines showed high levels of STAT3, an important transcription factor activated downstream of EGFR, and after EGF stimulation both showed phosphorylation of serine 727 (fig S3b) but not of tyrosine 705 (data not shown).

**Figure 4:**
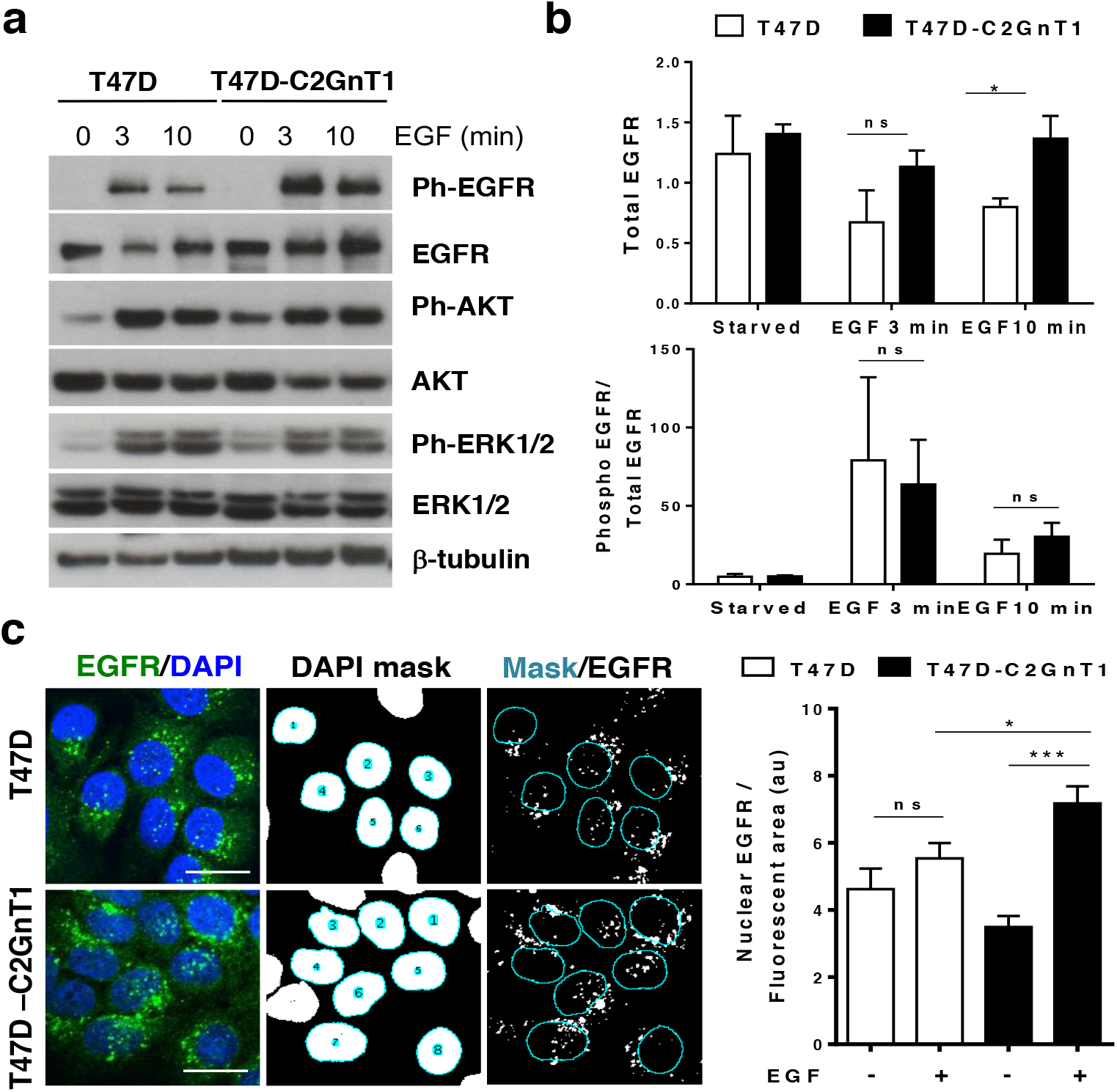
O-linked glycosylation does not affect EGF induced signaling but influences nuclear localisation. (a) T47D and T47D-C2GnT1-J cells were starved for 24 hours and treated with EGF (100ng/ml) for the indicated time. Cell extracts were subjected to western blotting using beta-tubulin, total EGFR, phospho-T1173 EGFR, AKT, phospho-S473 AKT, ERK1/2 and phospho-ERK antibodies. Representative western blots are shown. (b) Results are expressed as the densitometric ratio of total EGFR to beta-tubulin (top panel); phospho-T1173 EGFR to total EGFR (bottom panel) and normalized to beta-tubulin (*P<0.05, n.s. non-significant; analysis by t-student test n=3). (c) T47D and T47D-C2GnT1-J cells were treated with EGF (100ng/ml) for 20 min and fixed for immunostaining with an anti-EGFR antibody (green) and DAPI (blue). Confocal images of T47D and T47D-C2GnT1-J cells are shown: (Left panels) EGFR (green) and DAPI (blue) overlay. (Middle panels) Mask corresponding to the nuclear area as defined by DAPI staining. (Right panels) Overlay of DAPI mask onto EGFR image to select nuclear associated EGFR signal. Graph shows quantification of nuclear associated EGFR, n=200 cells from 3 independent experiments. (*P<0.05, ***P<0.001, n.s. non-significant; analysis by t-student test).

In addition to activation of signaling from the cell membrane, following endocytosis, EGFR can traffic to the nucleus and bind to the promoters of genes regulating transcription (15, 17, 22). Therefore, a change in the localization of EGFR could be a possible mechanism to explain the differential transcriptional program induced by EGF in these cells. Because N-glycosylation can affect receptor turnover and endocytosis (23), we studied the effect of O-glycosylation in the sub-cellular localization of EGFR by confocal microscopy. We observed the internalization of EGFR upon EGF binding in both cell lines (fig S3c) and in both cases we observed EGFR associated to endosomes as indicated by EGFR-CD63 co-localization (fig S3d). Quantification of nuclear associated endosomes, endosomes that overlap with nuclear staining in confocal images taken at the level of maximum cell diameter (24), showed that there is a higher of ERFR nuclear stain in T47D-C2GnT1-J cells compared to T47D cells (fig 4c). Subcellular fractionation studies showed that, although the level of EGFR in the nucleus was relatively low, EGFR was found in the nuclear fraction of T47D and T47D-C2GnT1-J and meanwhile EGFR declined in the membrane + cytoplasm fraction after stimulation with EGF in both cell lines, EGFR levels were maintained in the nuclear fraction of T47D-C2GnT-J cells (fig S3e).

Together these results show that upon EGF binding, in T47D-C2GnT1-J cells that carry branched core 2 glycans, a significant increase in EGFR nuclear associated endosomes is observed compared to T47D carrying core 1-based glycans.

### EGFR, GAL3 and MUC1 form a complex upon EGFR activation of T47D carrying core 1 O-glycans but not in T47D carrying core 2 O-glycans

Galectin-3 is a lectin that binds preferentially to terminal beta-galactosides on mature N-glycans and elongated core 2 O-glycans (25). Galectin-3 has been shown to bind to the extracellular domains of EGFR and MUC1 (26, 27). We immunoprecipitated EGFR from solubilized membrane and cytoplasmic cell extracts of T47D and T47D-C2GnT1-J cells and observed that EGFR forms a complex with galectin-3 and MUC1 in T47D cells treated with EGF but not in resting cells, as has been previously described (28). In contrast, upon EGF stimulation no MUC1/galectin-3/EGFR complex was observed in T47D-C2GnT1-J cells (fig 5a). Interestingly some interaction of EGFR with galectin-3 was observed in unstimulated T47D-C2GnT-J cells but this was at a very low level (fig 5a).

**Figure 5:**
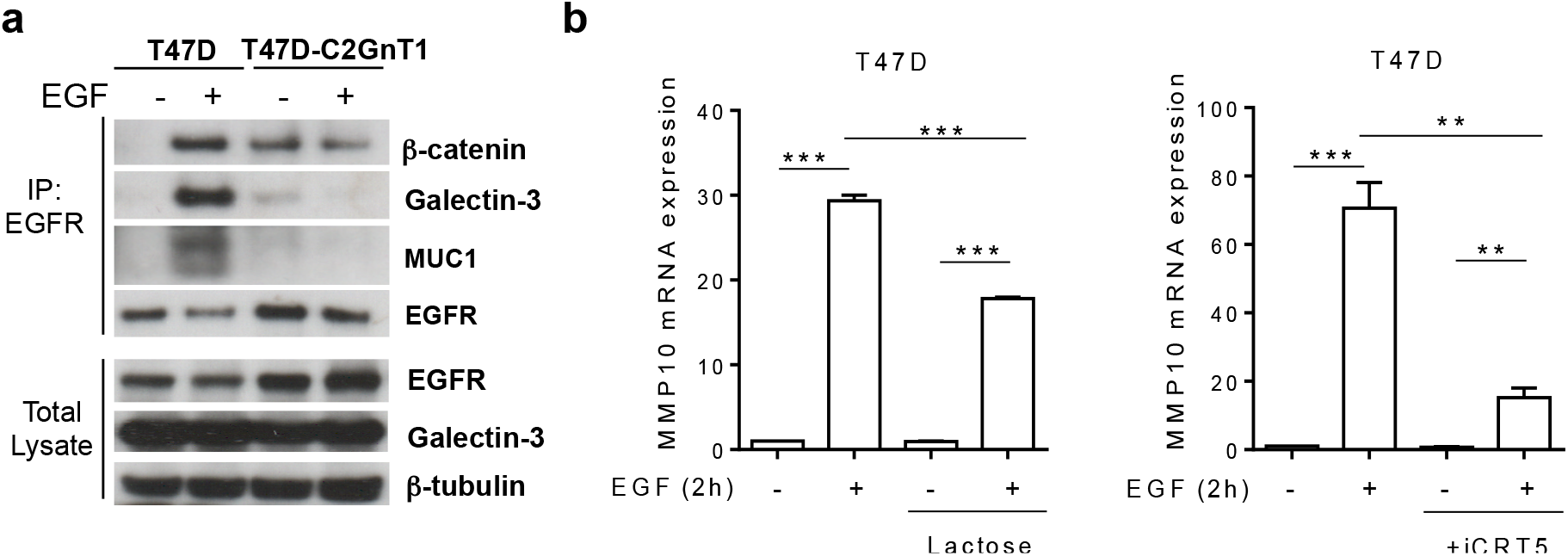
O-linked glycosylation dictates EGFR/galectin-3/MUC1 complex formation. (a) T47D or T47D-C2GnT1-J cells were starved for 24 hours and treated or not with EGF (100 ng/ml) for 3 min. Total cell extracts and EGFR immunoprecipitates (IP) were analysed by western blotting (IB) with the indicated antibodies. (b) T47D cells were starved for 24 hours, then pretreated with lactose (50mM), iCRT5 (50μM) or vehicle for 6 hours, followed by EGF (100 ng/ml) or nothing for 2h. Transcript levels of MMP10 were assessed by qPCR using PUM1 as a housekeeping gene and shown relative to the expression in starved, non-treated T47D cells (**P<0.05, ***P<0.001; analysis by t-student test n=4).

β-catenin, an important transcriptional regulator in breast cancer cells interacts with EGFR (29) and binds to the phosphorylated cytoplasmic tail of MUC1 (30). We therefore investigated if β-catenin was in a complex with EGFR and MUC1 in T47D cells. We observed that β-catenin precipitated with EGFR in EGF stimulated T47D cells but not in starved cells (fig 5a). However, in T47D-C2GnT-J cells we observed β-catenin to be in a complex with EGFR in non-stimulated and EGF stimulated cells in the absence of MUC1. These results show that extended core 2 O-linked glycosylation inhibits the interaction of EGFR with galectin-3 and MUC1 but facilitates the constitutive interaction of EGFR with β-catenin.

To study the relevance of galectin-3 binding and β-catenin interaction in the EGF induced transcriptional program of core 1 vs core 2 O-glycosylated cells, we looked at the expression of MMP10 in cells pretreated with lactose, which competes for binding of galectin-3, affecting only the extracellular interactions of galectin-3, or treated with iCRT-5, that inhibits β-catenin/LCF interaction, specifically blocking β-catenin transcriptional activity (31). We observed that pre-treatment with the β-catenin inhibitor or inhibition of galectin-3 binding significantly inhibited the upregulation of MMP10 in T47D cells stimulated with EGF (fig 5b). However none of these treatements affected the upregulation of CX3CL1 in T47D-C2GnT1-J cells stimulated with EGF (fig S4). Together these results suggest that the galectin-3 mediated formation of the complex β-catenin-EGFR-MUC1 is required for the specific transcriptional upregulation of MMP10 in T47D breast cancer cells.

## DISCUSSION

Changes in glycosylation are very common in cancer and although increased sialylation is a common event, different tumour types can show different glycosylation patterns. Indeed different cancers can be clustered according to the expression of glycosyltransferases (32). The two O-linked glyco-phenotypes investigated here represent core 1 and sialylated core 1-based glycans (T47D), and core 2-based glycans (T47D-C2GnT1). While sialylated core 1 glycans are found on the majority of breast cancer cells, and normal mammary epithelial cells express glycoproteins carrying exclusively core 2 O-linked glycans, the differential expression of these glycans is not absolute and in ER-ve breast cancers branched core 2 glycans appear to dominate although core 1 based glycans are also present (8).

Upon binding of EGF, EGFR dimerizes and activates its tyrosine kinase activity leading to a plethora of down-stream signalling pathways. However, it is now clear that ligand binding can also stimulate the nuclear localisation of EGFR (22, 33–35) resulting in its direct involvement in the regulation of transcription. Here we show for the first time that the O-linked glyco-phenotype of the cell can influence the pattern of gene transcription induced by EGF binding to EGFR. As changes in O-linked glycosylation are very common in the transition to malignancy this finding is highly significant to EGFR driven cancer progression.

Global gene analysis showed that many of the genes expressed in the two glyco-phenotypes encode secreted proteins and proteases involved in cell:cell and cell:extracellular matrix interactions. The product of these genes regulate the half-life and/or activation of secreted or membrane receptors that through autocrine and paracrine functions they orchestrate processes like angiogenesis, immunosupression and extracellular matrix remodeling (36, 37). MMP10, FGFBP1 and CXC3L1 have all been implicated in cancer and showed differential expression between the two phenotypes. MMP10 is significantly upregulated in ER+ve breast cancer that carry linear, core 1 glycans as seen in T47D cells. In contrast the chemokine CX3CL1 and FGF-BP1, were more highly expressed in the core 2 carrying glyco-phenotype and our analysis shows that expression in breast cancer is highly correlated with the ER-ve subtype which carries more core 2 O-linked glycans.

The lectin galectin-3 can exert multiple cellular functions (25) and binds to EGFR (26) and MUC1 (27). We demonstrated that different O-linked glycan cores carried by cells can influence the formation of cell surface complexes of EGFR with galectin-3, MUC1 and beta-catenin. EGFR expressed by cells carrying core 1 based O-linked glycans can interact with galectin-3 and MUC1, while EGFR expressed by cells carrying core 2 based glycans show no such interaction. As well as carrying O-linked glycans both EGFR and MUC1 have predicted N-glycosylation sites. Although the binding site for human galectin-3 on EGFR has not been identified, in mice, Mgat5 dependent N-glycosylation of EGFR is necessary for galectin-3 binding (23). Moreover, interaction of MUC1 with galectin-3 has also been shown to be through N-linked glycosylation (27) although others have shown that core 1 O-linked glycans may also be involved (38). Our data suggest that bulky core 2 O-glycans carried on the surface of glycoproteins may sterically impede the access and binding of galectin-3 to the N-glycans in MUC1 and EGFR. In contrast upon conformational changes induced by EGF the short core 1-based O-glycans present in T47D cells allow binding of galectin-3, possibly to the N-linked glycans on EGFR and MUC1. However, it cannot be ruled out that galectin-3 may bind directly to the core 1 O-linked glycans on MUC1 and EGFR in T47D cells (39).

In normal mammary epithelial cells MUC1 carries only branched core 2 glycans and is exclusively found on the luminal surface of the cells (40). In contrast, EGFR is found on the baso-lateral surface (41) and therefore in normal cells MUC1 and EGFR are spacially separated preventing interaction. However, in the change to malignancy cell polarity is lost and MUC1 is found all over the cell surface allowing interaction with EGFR which we have shown can be influenced by the glycosylation state of the cell.

Others have shown that MUC1 interacts with EGFR and modulates its stability by preventing its ubiquitination (15) and that MUC1 expression can contribute to EGF induced signaling pathways (39). However, we did not see any significance differences between cell lines with different O-glycosylation patterns in the downstream signaling from EGFR or the ubiquitination of EGFR upon EGF binding (data not shown). This suggests that the difference between the total levels of EGFR observed between T47D and T47D-C2GnT1 may be due to differential trafficking/recycling of the receptor and indeed we found EGFR accumulation in the nuclear endosomes and nuclear extracts in core 2 carrying than in the core 1 carrying T47D cells.

Our data suggest that O-glycosylation can influence EGFR intracellular trafficking and the formation of EGFR complexes at the plasma membrane upon EGF stimulation in breast cancer cells. We showed that inhibition of the action or formation of these complexes can inhibit specific gene transcription. The fact that aberrant O-linked glycosylation is such a common and consistent change in cancers indicates that it plays an important role in tumor growth and progression. The data presented here demonstrates a further functional outcome as a result of changes in O-glycosylation in cancer and illustrates the crucial role of glycans play in malignancy including influencing the transcriptional programme.

## MATERIALS AND METHODS

### Cell culture and transfection

T47D were obtained from their originator; Dr Keydar and were cultured in RPMI-1640 medium (Lonza) supplemented with 10% FCS (Life technologies). All media were supplemented with 2 mM L-glutamine, 100 U/ml penicillin and 100 µg/ml streptomycin. T47D cells transfected with *GCNT1* as described in [1] were additionally cultured with 500 μg/ml G418. 100ng/ml EGF, 50 μM iCRT5 and 50 mM lactose were used alone or in combination to stimulate cells for the times indicated. Master and working banks were made of all cell lines and cells kept in culture no longer than 8-10 weeks. ZR-75-1 and BT20 were authenticated by LGC Standards in February 2011 using the short-tandem repeat profile of 16 loci. These contain the 9 loci used by the American Type Culture Collection and both lines exactly matched these profiles

### Cell lysis, cell fractionation and western blotting

Cells were lysed after EGF stimulation in lysis buffer (50 mM Tris-HCl pH 8, 0.5 mM EDTA, 150 mM NaCl, 1% NP-40 supplemented with protease and phosphatase inhibitor cocktails (Roche). Lysates were homogenised through a needle and centrifuged for 10 min at 16,000g to separate soluble from insoluble fractions. Protein lysates were analysed by SDS-PAGE and immunoblotting (Invitrogen). Bound antibodies were visualised with horseradish peroxidase-conjugated anti-IgG antibodies and enhanced chemiluminescence susbtrates (GE).

Cells were scraped, washed with PBS, resuspended in hypotonic buffer (10 mM Hepes (pH 7.9), 1.5 mM MgCl2, 10 mM KCl, 0.2 mM phenylmethylsulfonyl fluoride and 0.5 mM dithiothreitol), and allowed to swell on ice for 10 min. 1% NP40 was added and the nuclei in the homogenized lysates separated the from soluble cytoplasmic/membrane extracted by spinning at 3300g for 5 min at 4 °C. The nuclear pellet was resuspended in cold nuclear extraction buffer (20 mM Hepes pH=7.9, 100mM NaCl, 1.5 mM MgCl2, 1% Triton, 1 mM EDTA, 1 mM EGTA, 10% glycerol, 0.5 % deoxycholate, 0.1% SDS) for 30 min and centrifuged at 12,000g for 30 min. The supernatant was used as nuclear extract.

Soluble nuclear and cytoplasmic/membrane extracts were resolved by SDS-PAGE and western blot analysis. Antibodies were obtained from the following sources: anti beta-tubulin, CCND1, EGFR clone D38B1, phosphoS1173-EGFR, ERK1/2, phosphoERK1/2, phosphoS473 AKT, STAT3, phosphoS727 STAT3 (Cell Signalling); anti CT-MUC1 (CT1, homemade); AKT, Lamin B1 (Santa cruz), anti-Galectin-3 (eBioscience); ANGPTL4, MMP10 and FGF-BP1 (R&D systems-Biotechne); HRP-labelled anti-mouse, anti-rat and anti-rabbit antibodies (Dako).

### Cell staining, confocal microscopy and EGFR nuclear endosomes quantification

Cells seeded on glass coverslips were fixed with 4% paraformaldehyde in PBS for 20 min, permeabilized with 0.1% Triton X-100 in PBS for 5 min and blocked with 5% BSA in PBS for a further 30 min. Cells were incubated for 1 h with anti-EGFR (cell signalling), anti-CD63 (Developmental Studies Hybridoma Bank (DSHB) and 4’,6-diamidino-2-phenylindole (DAPI) to stain nuclei. Alexa-fluor 488 or 546 secondary antibodies (Molecular Probes) were used. Images were generated with an A1 Nikon Eclipse-Ti-E Inverted confocal microscope using a 40x objective equipped with a DS-U3 Digital Sight Color Camera and NIS software. For quantification of EGFR-positive nuclear-associated endosomes images of cells stained with anti-EGFR and DAPI were analysed. ImageJ was used to analyse at least 100 cells per condition and to calculate EGFR positive area within the nucleus, delimited by a mask obtained using a thresholded DAPI staining and superimposed on the EGFR image (Nuclear EGFR). Same threshold was applied to all images.

### Flow cytometry

2×10^5^ cells were resuspended in PBS + 0.5% BSA and incubated with either anti-EGFR (42), HMFG1 or SM3 (8, 18) antibodies for 1h at 4°C then washed and incubated with FITC-conjugated anti-mouse, anti-rat or anti-rabbit antibodies. Isotype controls with only secondary antibodies were used.

### RNA isolation and qPCR

RNA was isolated from cells using RNeasy Mini-kits (Qiagen) and contaminating DNA digested on-column. For cDNA synthesis, a SuperScript VILO kit was used (Invitrogen). Quantitative real-time PCR (qPCR) was carried out with cDNA using SYBR green-containing mastermix (Primer Design) using PUM1 as a reference gene. Three technical replicates were used for each condition in each qPCR assay. See oligonucleotides used in this work in table S4.

### Genome wide expression analysis

Genome wide expression analysis was performed on Illumina Human HT12 v4 BeadArray according to the manufacturer guidelines. The data have been deposited in NCBIs Gene Expression Omnibus (GEO) and are accessible through GEO Series accession number GSE107629. Microarrays were normalized by using quantile normalization and probes with multiple genomic matches, or matches to non-transcribed genomic regions were discarded. Differentially expressed genes were measured using the moderated t-statistics with empirical Bayes shrinkage of the sample variances method from limma package as implemented in Bioconductor (43). P-values were corrected using false discovery rate (FDR) and only genes with an FDR less than 0.01 were used for further analyses. Gene Ontology Enrichment Analysis (GOEA) was performed using the Database for Annotation Visualization and Integrated Discovery (DAVID) (44). Heatmaps and volcano plots were generated using ggplot2 package as implemented in Bioconductor (45).

### Analysis of gene expression in breast cancers

Total RNA was extracted from ER-negative and ER-positive breast cancers and submitted to microarray analysis on the GlycoV4 oligonucleotide array, a custom Affymetrix GeneChip (Affymetrix, Santa Clara, CA) designed for the Consortium for Functional Glycomics as described in Julien *et al.* (8). Data is deposited in GEO with accession number GSE32394.

The METABRIC dataset, consisting of over 2,500 breast cancer samples, was interrogated for gene expression from cBioportal, http://www.cbioportal.org. The TCGA mRNA expression dataset deposited in Oncomine, https://www.oncomine.org for 76 invasive breast cancer samples and 61 normal breast tissue samples was interrogated for MMP10 expression.

## Statistical analysis

All data are presented as mean+SEM. The number of biological replicates (independent experiments in cell based assays) is stated in each figure legend as n. Statistical analysis was performed using GraphPad Prism software and statistical significance calculated using an unpaired two-tailed t-test (for comparing two conditions) or Mann-Whitney U-test (for expression analysis with METABRIC data). For the genome expression array a sample size of n=3 was used. Using statmate we determined that a sample size of 3 in each group has a 90% power to detect a difference between means of 50.51% half or double the expression) with a significance level (alpha) of 0.05 (two-tailed) using an unpaired *t* test.

## ACKNOWLEDGMENTS

This work was supported by grants from the Medical Research Council, MR/J007196/1 and MR/R000026/1. We thank the Nikon Imaging Center@King’s (John Harris, Dan Matthews and Isma Ali) for help with microscopy. We also thank Suzanne Papp and Lana Schaffer from the Consortium for Functional Glycomics, The Scripps Institute, La Jolla, for carrying out the glycan-related gene array analysis on the primary breast cancer samples and Dr Jelmar Quist with assistance with the METABRIC analysis.

## Supplementary File 1A

**Suppl. Figure 1.**
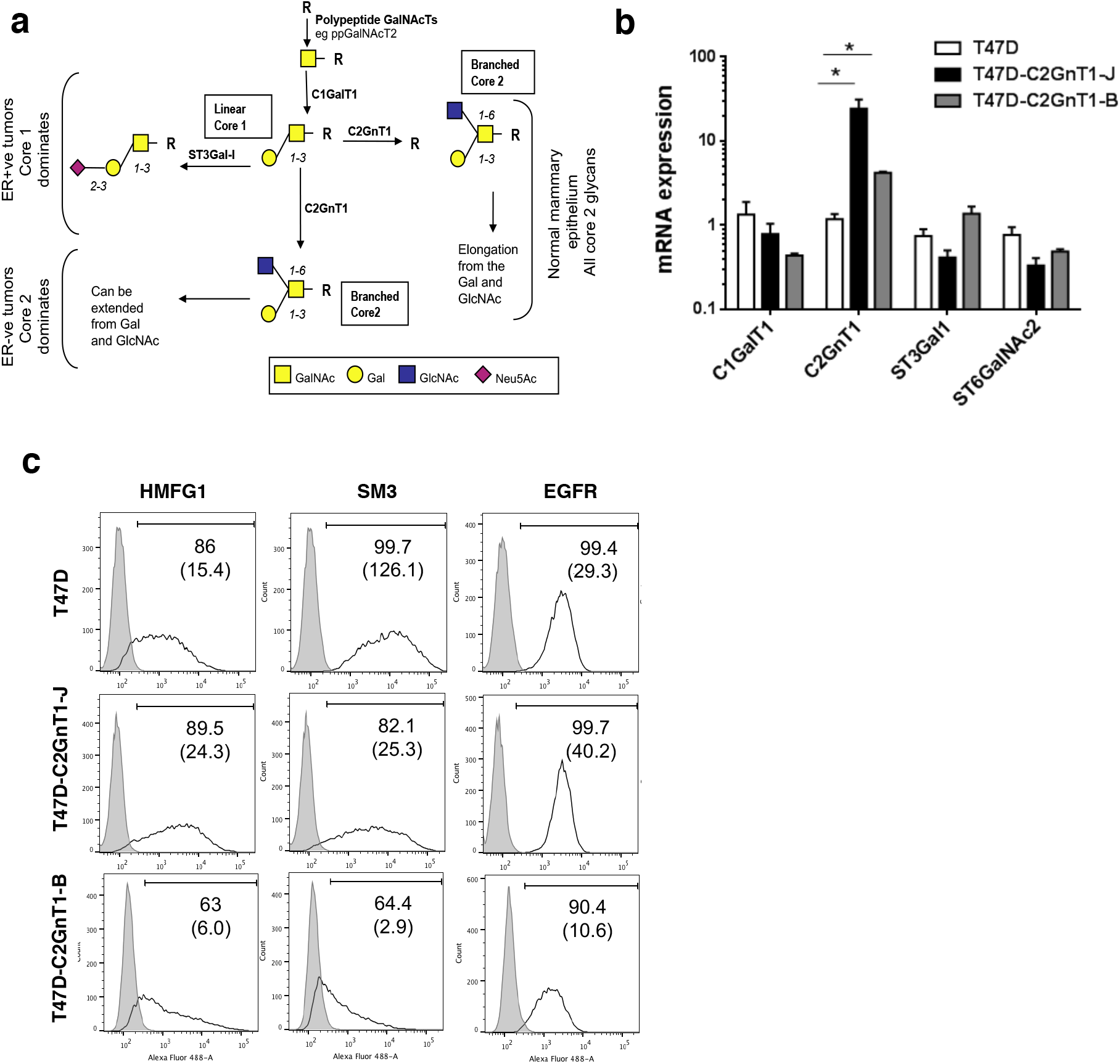
T47D and T47D-C2GnT1-J cells express similar levels of EGFR at the plasma membrane. (a) Pathways of O-glycosylation in breast tissue and tumors. In normal mammary epithelial cells, core 1 glycans are converted to branched core 2 by the action C2GnT1. In breast carcinomas this conversion is reduced mainly due to overexpression of ST3Gal-I sialyltransferase resulting in the expression of mainly core 1-based structures which are largely sialylated. However, glycosyltransferase mRNA expression analysis suggest this is associated with ER+ breast cancers and in ER-ve tumours more branched core 2 glycans can be found. (b) C1GalT1, C2GnT1, ST3Gal1 and ST6GalNAc2 glycosyltransferases transcript levels were assessed in T47D, T47D-C2GnT1-J and T47D-C2GnT1-B cells by quantitative PCR using PUM1 as a housekeeping gene, quantified using the ΔΔCt method and shown relative to the levels present in T47D cells (# P<0.05 analysis by t-student test (n=3). (c) MUC1 (HMFG1), short core 1 O-glycan MUC1 (SM3) and EGFR expression in T47D and T47D-C2GnT1-J and -B clones. T47D, T47D-C2GnT1-J and T47D-C2GnT1-B cells were stained with HMFG1, SM3, EGFR or isotype antibodies (grey). Binding populations were visualized using flow cytometry. Figures represent percentage of interacting cells and median fluorescence intensity in brackets (n=3).

**Suppl. Figure 2:**
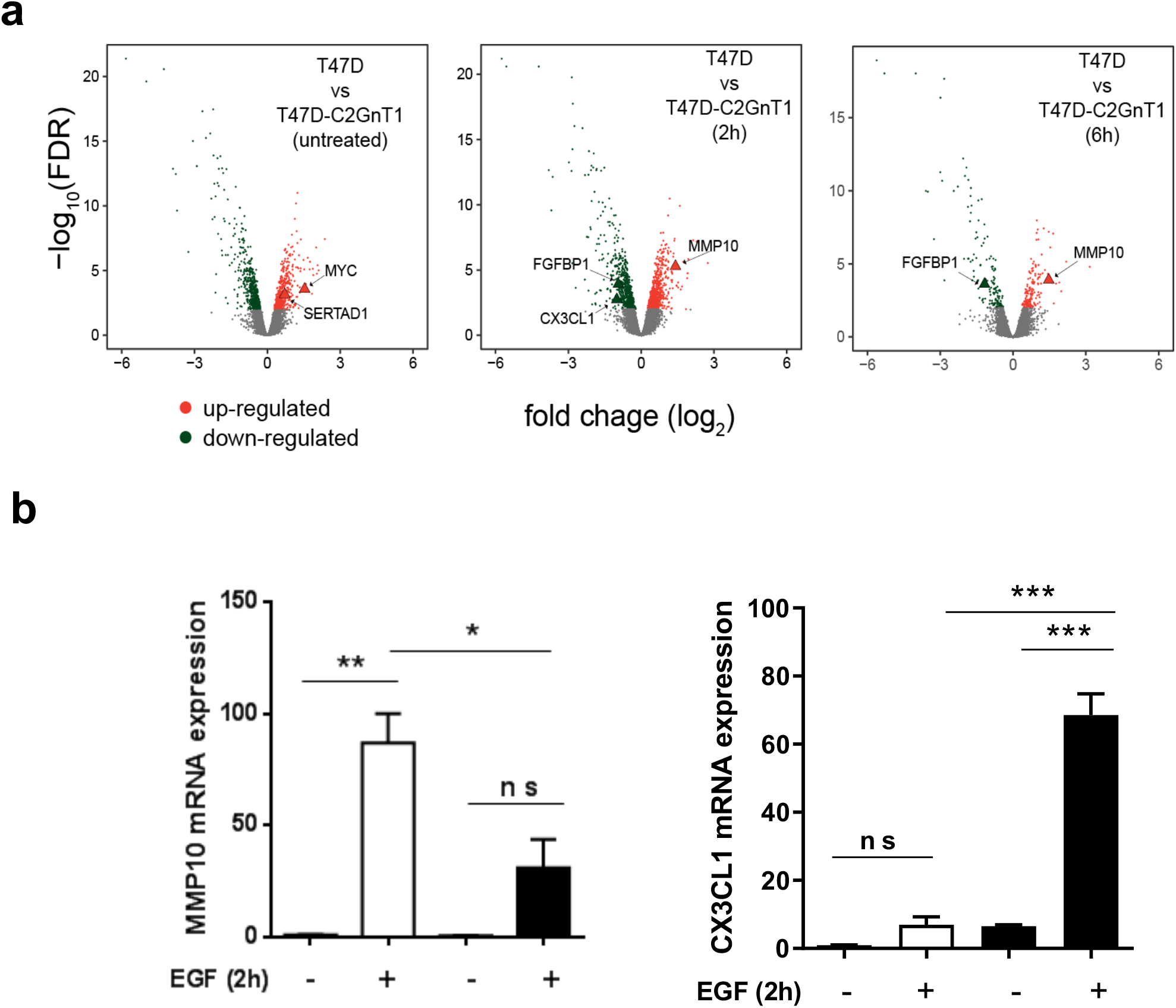
Differential gene expression between two glyco-phenotypes. (a) T47D and T47D-C2GnT1-J cells were treated as in figure 1c of main text. Volcano plots of differentially expressed genes between T47D core-1 and T47D core-2 breast cancer cells before and after 2h or 6h of treatment with EGF are shown. Significantly differentially expressed genes (FDR < 0.01) in cells before treatment (untreated) or after 2h or 6h of treatment with EGF 100ng/ml are highlighted in red (up-regulated) and green (down-regulated). Relevant transcriptional targets of EGF signaling are highlighted as triangles. (b) T47D and a second clone of T47D-C2GnT1-B were starved for 24 hours and treated with EGF (100ng/ml) for 2h. Transcript levels of MMP10 and CX3CL1 were assessed by quantitative PCR using PUM1 as a housekeeping gene, quantified using the DDCt method and shown relative to the expression in starved, non-treated T47D cells (*P<0.05, **P<0.01; analysis by t-student test n=3).

**Suppl. Figure 3:**
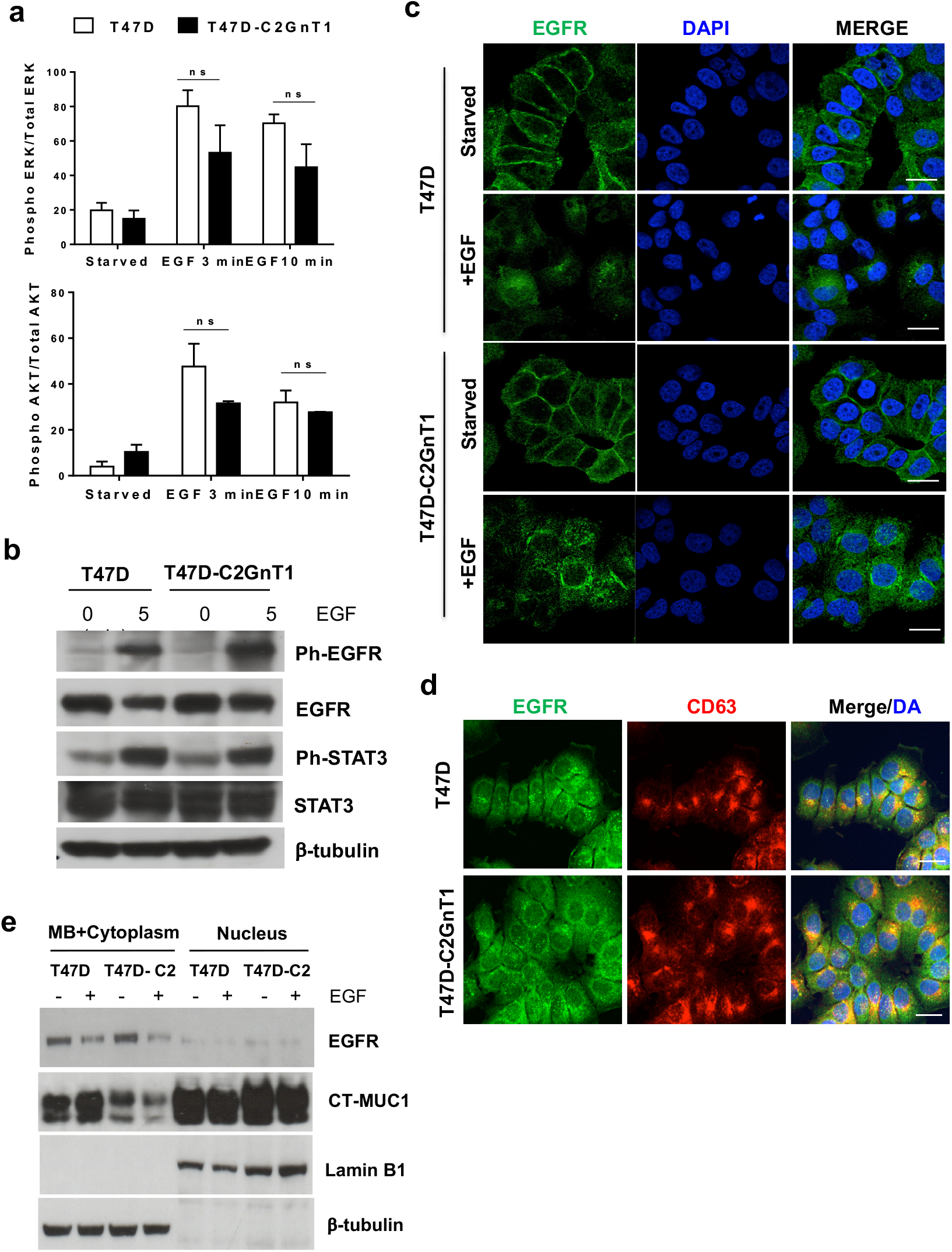
O-linked glycosylation does not affect EGF induced signaling but influences nuclear localization. (a) T47D and T47D-C2GnT1-J cells were starved for 24 hours and treated with EGF (100ng/ml) for the indicated time. Quantification of ERK and AKT activation from western blots shown in figure 4a, main text. (b) Cells were treated as in (a) and lysates analysed by immunoblotting with the indicated antibodies. (c) T47D and T47D-C2GnT1-J cells were starved for 24 hours and treated or not with EGF (100ng/ml) for 20 min. Cell were fixed for immunostaining to visualize EGFR and the nucleus (DAPI). Confocal images of the central region of T47D and T47D-C2GnT1-J cells stimulated or not with EGF (100ng/ml) are shown. (d) T47D and T47D-C2GnT1-J cells treated as described before (d) were fixed for immunostaining to visualize EGFR, late endosomes (CD63) and the nucleus (DAPI). Confocal images of EGF treated cells are shown. (e) Cells were starved for 24 hours and treated or not treated with EGF (100ng/ml) for 20 min. Cell extracts were fractionated into membrane plus cytoplasm and soluble nuclear fractions and subjected to western blotting using beta-tubulin, Lamin B1, total EGFR and CT-MUC1 antibodies. Representative western blots are shown (n=2).

**Suppl. Figure 4:**
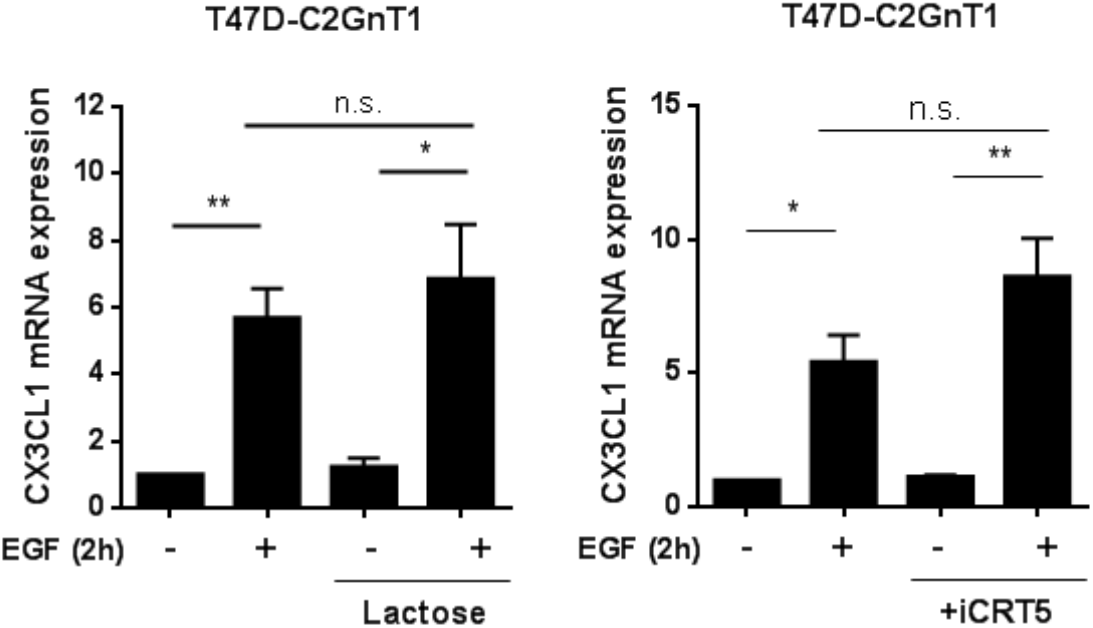
Inhibition of galectin/ MUC1/EGFR complex and β-catenin inhibition does not affect expression of CXC3CL1. T47D cells were starved for 24 hours and then pretreated with lactose (50mM), iCRT5 (50μM) or vehicle for 6 hours, followed by EGF (100 ng/ml) or nothing for 2h. Transcript levels of CX3CL1 were assessed by quantitative PCR using PUM1 as a housekeeping gene, quantified using the ΔΔCt method and shown relative to the expression in starved, non-treated T47D cells (**P<0.05; analysis by t-student test n=4).

## Supplementary File 1B

**Table S3:**
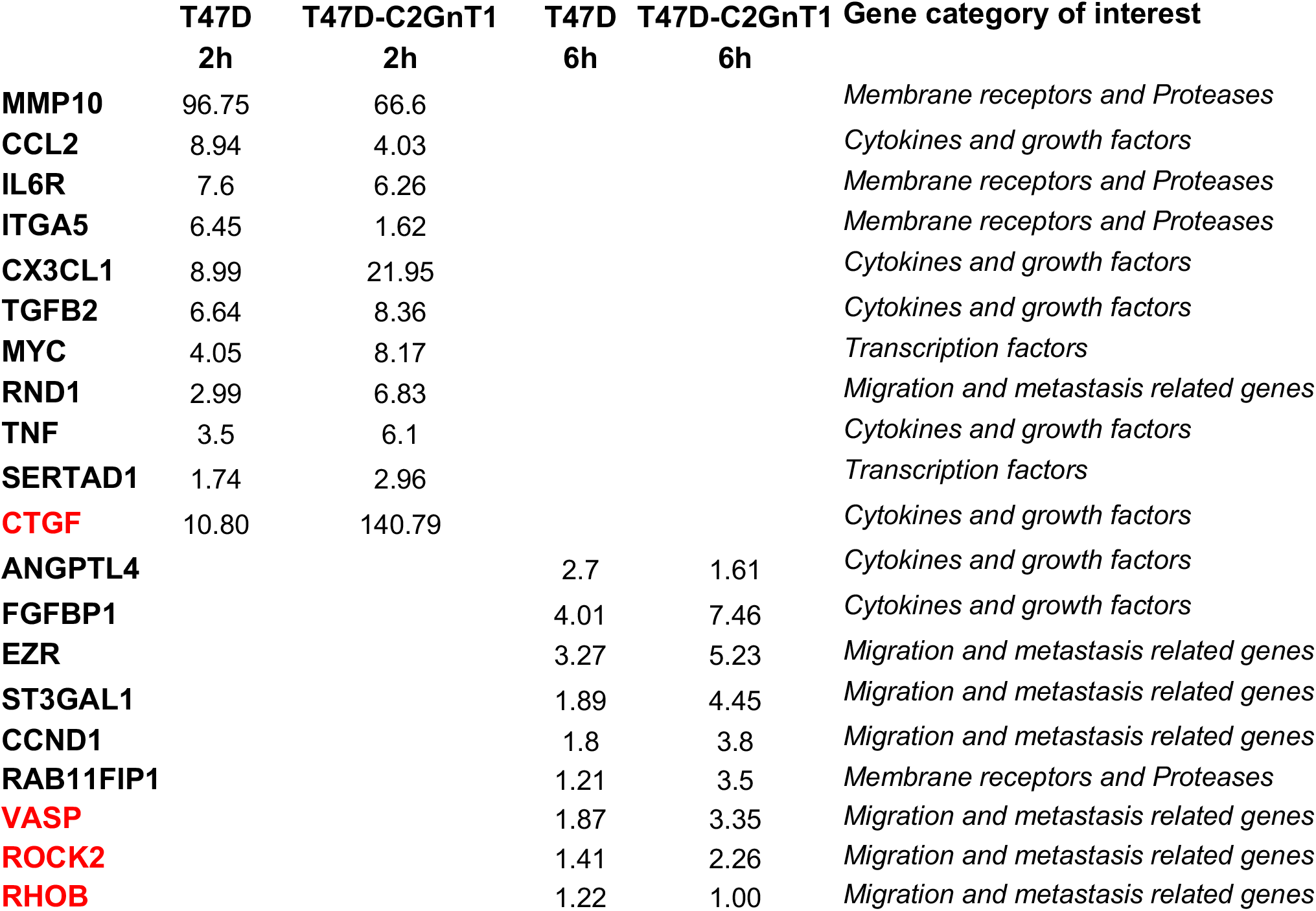
Summary of fold change in gene expression upon EGF stimulation for the 20 genes selected for validation. The table shows the of average fold change in mRNA expression from stimulated to non-stimulated conditions assessed by quantitative PCR from three independent experiments and the gene category of interest. PUM1 was used as a housekeeping gene. mRNA levels were quantified using the ◻◻Ct method. In red are those that showed no significant upregulation in any of the glyco-phenotype cell lines or showed different expression to that observed in the microarray.

